# Modulation of fear generalization by the zona incerta

**DOI:** 10.1101/485250

**Authors:** Archana Venkataraman, Natalia Brody, Preethi Reddi, Jidong Guo, Donald Rainnie, Brian George Dias

## Abstract

Fear expressed towards threat-associated stimuli is an adaptive behavioral response. In contrast, the generalization of fear responses toward non-threatening cues is maladaptive and a debilitating dimension of trauma- and anxiety-related disorders. Expressing fear to appropriate stimuli and suppressing fear generalization requires integration of relevant sensory information and motor output. While thalamic and sub-thalamic brain regions play important roles in sensorimotor integration, very little is known about the contribution of these regions to the phenomenon of fear generalization. In this study, we sought to determine whether fear generalization could be modulated by the zona incerta (ZI), a sub-thalamic brain region that influences sensory discrimination, defensive responses, and retrieval of fear memories. To do so, we combined differential intensity-based auditory fear conditioning protocols in mice with C-FOS immunohistochemistry and DREADD-based manipulation of neuronal activity in the ZI. C-FOS immunohistochemistry revealed an inverse relationship between ZI activation and fear generalization – the ZI was less active in animals that generalized fear. In agreement with this relationship, chemogenetic inhibition of the ZI resulted in fear generalization, while chemogenetic activation of the ZI suppressed fear generalization. Furthermore, targeted stimulation of GABAergic cells in the ZI reduced fear generalization. To conclude, our data suggest that stimulation of the ZI could be used to treat fear generalization in the context of trauma- and anxiety-related disorders.

## INTRODUCTION

Expressing fear toward cues that had been previously associated with trauma is adaptive (conditioned fear). Equally adaptive is the expression of fear toward stimuli that closely resemble traumatic cues (generalization). Such generalization of fear allows the organism to be “better safe than sorry”. However, fear generalization can diminish quality of life and is a highly debilitating dimension of trauma- and anxiety-related disorders like Post-Traumatic Stress Disorder (PTSD) and Generalized Anxiety Disorder (GAD) (1-4). Reducing fear generalization while maintaining adaptive fear responses will reduce the daily burden experienced by individuals living with these disorders. Recently, introducing procedures that involve stimulus discrimination into cognitive behavioral therapy has been shown to reduce fear generalization, re-experiencing and intrusive thoughts in PTSD patients (5-7).

Brain regions such as the lateral amygdala (8-10), central amygdala (11, 12), prefrontal cortex (13, 14), hippocampus (3, 15), and bed nucleus of the stria terminalis (BNST) (16, 17) have been implicated in fear generalization. More importantly, these regions play crucial roles in detecting threats and assigning valence to environmental stimuli (18-21). Therefore, while manipulating these regions could potentially reduce fear generalization, doing so might compromise threat detection, conditioned fear and survival. In this study, we set out to ask whether targeting brain regions outside of the aforementioned canonical fear-related circuitry could reduce fear generalization.

Thalamic and sub-thalamic brain regions are ideal candidates to exert modulatory control over appropriate fear expression because they serve as hubs relaying information from sensory cortices to limbic, midbrain and brainstem nuclei (15, 22-24). The contributions of these brain regions have mostly been ignored in the context of fear-related behavior. At the level of the thalamus, studies have demonstrated that the auditory thalamus influences fear generalization (25, 26) and that the paraventricular thalamus influences fear conditioning and fear memory retrieval (27, 28). Most recently, the zona incerta (ZI), a sub-thalamic region, has received attention for its role in modulating defensive responses and retrieval of fear-related memories (29, 30). Notably, studies in rodents have highlighted that the ZI influences sensory discrimination (31, 32) and that stimulation of the ZI in humans facilitates discrimination of fearful from non-fearful stimuli (33). Motivated by these findings, we hypothesized a potential role for the ZI in fear generalization.

To test this hypothesis, we leveraged the fact that high threat intensities elicit excessive fear responses even towards neutral stimuli – fear generalization. We used differential auditory fear conditioning in mice at varying threat intensities to model high and low threat conditions. More specifically, during conditioning, auditory conditioned stimulus (CS+) presentations were paired with foot-shocks of low (0.3mA) or high (0.8mA) intensity, whereas a second stimulus (CS-) was not reinforced. Animals trained under low threat conditions (0.3mA) expressed appropriately high fear responses to CS+ and relatively low fear responses to CS-. However, animals trained under high threat conditions (0.8mA) expressed high fear responses to both CS+ and CS-, exhibiting maladaptive fear generalization as is observed in individuals affected by PTSD and GAD. C-FOS immunohistochemistry revealed that the ZI was less active in animals that exhibited fear generalization following training under high threat conditions.

To directly test whether the ZI plays a role in fear generalization, we manipulated cellular activity in the ZI using chemogenetic approaches. Decreasing cellular activity in the ZI resulted in fear generalization in animals trained under low threat conditions, while stimulating cells in the ZI suppressed fear generalization in animals trained under high threat conditions. With GABAergic cells in the ZI implicated in fear-related behaviors, we next asked whether stimulating GABAergic cells in the ZI would prevent fear generalization. Indeed, we found that fear generalization was reduced after targeted chemogenetic stimulation of GABAergic cells in the ZI of animals trained under high threat conditions. Notably, chemogenetic stimulation enhanced discrimination between the CS- and CS+, allowing for continued expression of fear toward the CS+. These results provide evidence that the ZI can modulate expression of appropriate behavioral fear responses. To our knowledge, our study is the first demonstration that stimulating the ZI may be of therapeutic value in reducing fear generalization.

## MATERIALS AND METHODS

### Animals

Adult female or male mice (2-3 months of age) were group-housed under a 14:10 light/dark cycle with food and water available ad libitum. C57BL/6J (wild type) mice and vGAT-CRE mice were originally ordered from Jackson labs and then bred in our vivarium for these experiments. All experimental procedures involving animals were approved by the Emory Institutional Animal Care and Use Committee and carried out in accordance with National Institute of Health standards.

### Auditory fear conditioning to test fear generalization

Differential intensity-based auditory fear conditioning was used to test fear generalization as described elsewhere (34). Briefly, the training and testing protocol consisted of four phases on four consecutive days: (1) habituation, (2) baseline, (3) training, and (4) testing (as outlined in Fig. 1A). On the first day, mice were habituated to the CS+ tone in the training context (Context A) for 10 minutes. One day later, during the baseline phase in Context A, freezing levels were measured during two random presentations each of the CS+ and the CS-, followed by exposure to continuous CS+ tone for a total of 10 minutes in Context A. Pre-exposure to the tones were designed in the protocol to allow for better discrimination and has been shown to prevent generalization (35, 36). The training phase that occurred one day later, included an initial 5-minute exposure to Context A followed by 20 trials consisting of 10 CS+ presentations that co-terminated with a 0.5 sec foot-shock with randomly interleaved 10 CS-presentations that were not reinforced. Depending on the experiment, either 0.3 mA (low threat condition) or 0.8 mA (high threat condition) foot-shocks were used as the unconditioned stimulus paired with the CS+. The inter-trial intervals varied randomly between 2-6 mins. During the testing phase on day 4, mice were exposed to a new context (Context B) for 3 minutes followed by two randomized presentations each of the CS+ and CS- and freezing levels measured during the tone presentations were used as a behavioral index of fear generalization. FreezeFrame-4 software (Actimetrics) was used for stimulus presentations and video recording of freezing behavior. Hardware associated with these experiments was purchased from Harvard Apparatus. The time spent freezing to CS+ and CS-was analyzed by an experimenter blind to the treatment condition, using FreezeFrame software with the freezing bout length set to 0.5 secs. Context A consisted of grid flooring, illuminated with house lights and cleaned with the disinfectant, quatricide. Context B consisted of plexiglass flooring, illuminated with infra-red lights and cleaned with 70% ethanol. Sound levels were adjusted so that all tones were presented at approximately 85dB. CNO injections (where relevant) were administered intra-peritoneally at a dose of 1 mg/kg and one hour before testing for fear generalization in Context B. Discrimination index (DI) was calculated as the difference in the % of time spent freezing to the conditioned and neutral tone divided by the sum of the % of time spent freezing to both tones.

**Figure 1:**
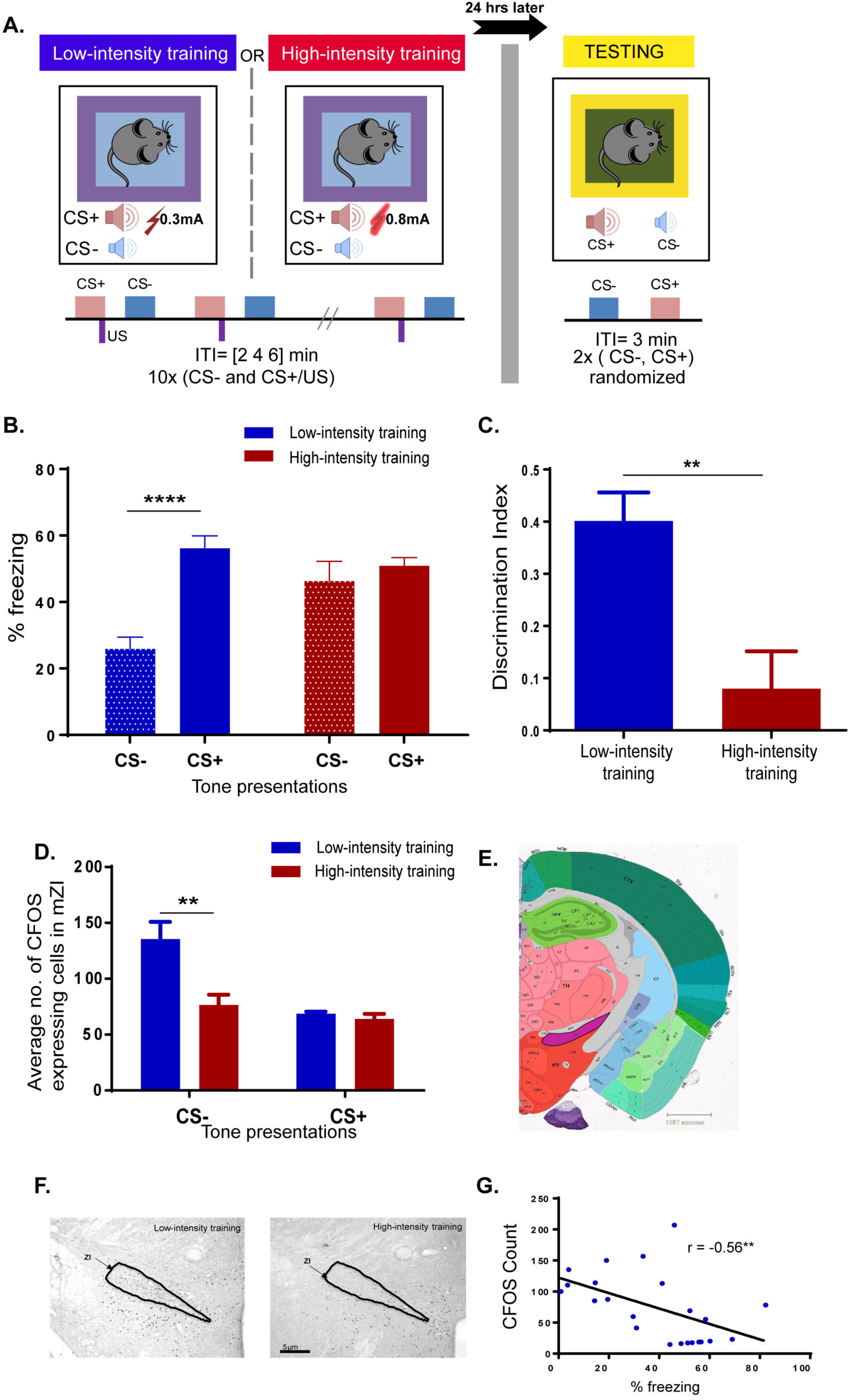
Fear generalization is associated with decreased neuronal activation in the ZI. **(A)** Outline of the differential auditory fear conditioning protocol used in the study. On day 1, one group of mice received CS+ tone presentations paired with 0.3mA foot-shocks (low threat intensity) and unpaired CS-tone presentations. Another group of mice received CS+ tone presentations paired with 0.8mA foot-shocks (high threat intensity) and unpaired CS-tone presentations. On day 2, freezing responses in both groups of animals were recorded for the CS+ and CS-tone presentations. **(B)** Animals trained under low threat conditions show low freezing response to CS- and high freezing response to CS+ (no fear generalization). In contrast, animals trained under high threat conditions show increased freezing response to both CS- and CS+ (fear generalization). **(C)** Discrimination indices calculated for the two groups reveal significant fear generalization in the animals trained under high threat conditions. **(D)** Decreased C-FOS expression was observed in the ZI of animals that showed increased fear to CS-presentation on testing day. **(E)** Reference image from Allen Brain Atlas showing position of ZI shaded in purple. **(F)** Representative images of C-FOS expression in the ZI in response to tone presentations during testing day after training under low or high threat conditions. **(G)** Significant correlation was found between C-FOS expression in the ZI and behavioral fear responses. \**p*<0.05, \*\**p*<0.01. \*\**p*<0.01, \*\*\*\**p*<0.0001. Data represented as Mean ± S.E.M.

### Stereotaxic surgeries

To manipulate cellular activity in the ZI of wild-type C57BL/6J animals we used AAV5-hSyn-hM4DGi-mCherry (to reduce activity), AAV5-hSyn-hM3DGq-mCherry (to stimulate activity) and AAV5-hSyn-GFP (as control) viruses. To stimulate cellular activity in the ZI of vGAT-CRE animals, we used AAV5-hSyn-DIO-hM3DGq-mCherry (to stimulate activity) and AAV5-hSyn-DIO-mCherry (as control) viruses. All viruses were obtained from the UNC Viral Vector Core and Addgene. Bilateral stereotaxic AAV injections into ZI were performed while the animal was under anesthesia using the following stereotaxic co-ordinates: AP: −1.52 mm, ML: 0.73 mm and DV: −4.79 mm relative to Bregma. AAV-containing solutions were injected at the rate of 1 nl/sec using Nanoject III (Drummond Scientific) and experiments were performed after 2 weeks to allow for optimal viral expression. A final volume of 50 nl of AAV5-hSyn-GFP, AAV5-hSyn-hM4DGi-mCherry, and AAV5-hSyn-hM3DGq-mCherry and 80 nl of AAV5-hSyn-DIO-mCherry or AAV5-hSyn-DIO-hM3DGq-mCherry was infused.

### C-FOS immunohistochemistry & Cell Counting

C-FOS protein expression was detected 90 minutes after exposure to either the CS+ or the CS-on testing day (as outlined in Supplementary Fig. 2A) in animals trained under low or high threat conditions. Mice were trans-cardially perfused with 4% paraformaldehyde dissolved in phosphate-buffered saline (PBS). Brains were removed and stored in paraformaldehyde solution for a day and transferred to 30% sucrose solution for 3-4 days before sectioning on a freezing microtome (Leica). 35μm brain sections were washed three times in 1X PBS for 10 minutes and incubated in 0.3% hydrogen peroxide to block endogenous peroxidase activity. Sections were blocked in 1X PBS with 5% normal goat serum for 1 hour at room temperature and then incubated in primary rabbit polyclonal anti-C-FOS antibody (1:6000 dilution, Millipore ABE 457) overnight on a shaker at room temperature. The next day, sections were washed three times in 1X PBS for 10 minutes and then incubated in secondary biotinylated goat anti-rabbit IgG antibody (1:1000 dilution, Vector Laboratories BA-1000) for 2 hours. Following this, sections were treated for 1 hour with avidin-biotin peroxidase system (Vectastain Elite ABC kit, PK-6100) and visualized using 3,3’-diaminobenzidine (Sigma-Aldrich). Sections were mounted on SuperFrost Plus slides (Fisher Scientific) and after drying, slides were coverslipped using Permount (Fisher Scientific). Images of the ZI were captured using Nikon E800 microscope at 4X magnification and C-FOS expression quantified using MCID Core Imaging software. C-FOS immunoreactivity was quantified across three consecutive sections per animal in both left and right hemispheres.

**Figure 2:**
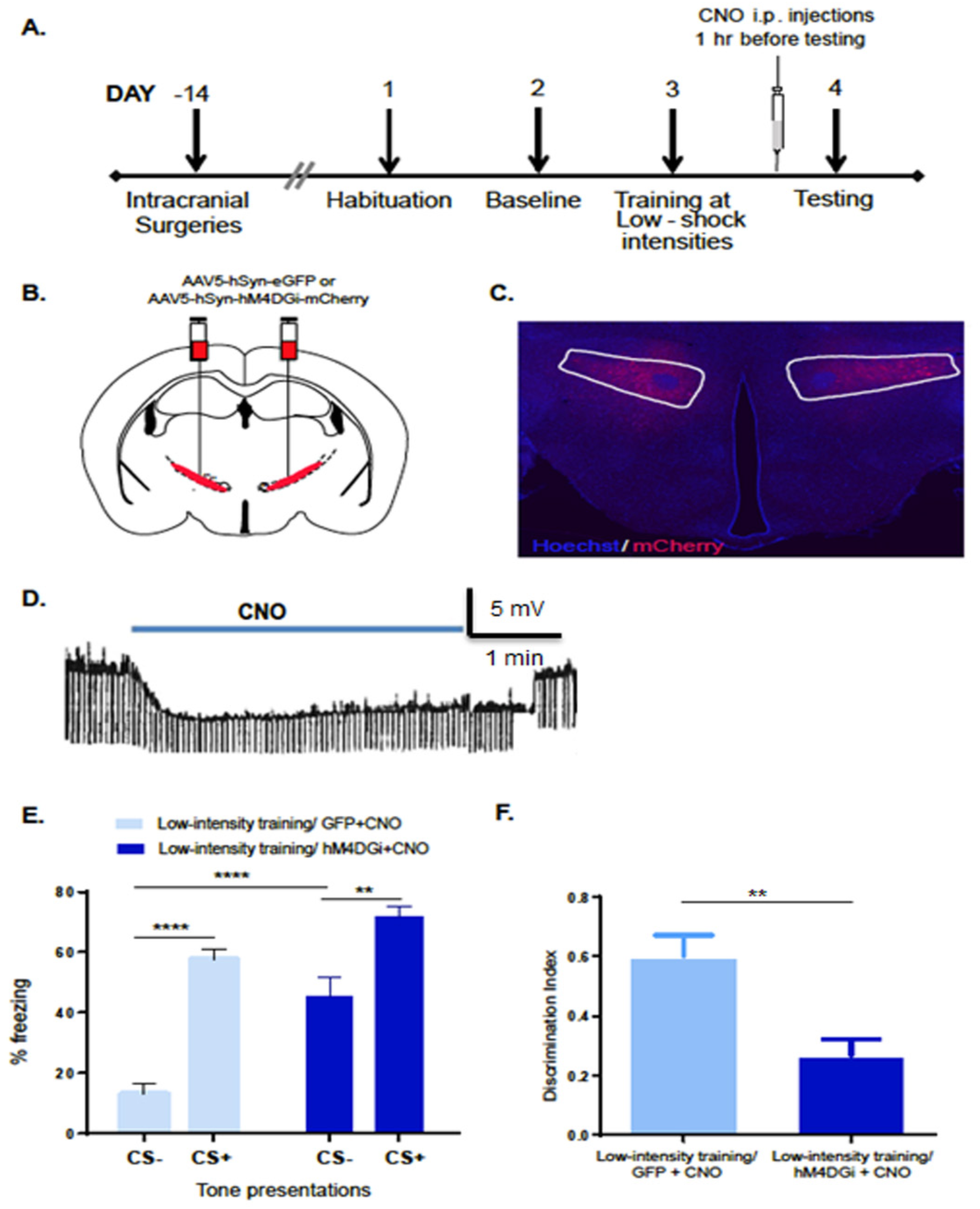
Decreasing cellular activity in the ZI results in fear generalization. **(A)** Experimental protocol for chemogenetic inhibition. Two weeks after intracranial injection of the control or DREADD virus, animals were conditioned to low threat intensities. The next day, CNO was administered intraperitoneally 1 hour before testing fear generalization. **(B)** Wild-type animals were injected with either the control virus (AAV5-hSyn-eGFP) or inhibitory DREADDs (AAV5-hSyn-hM4DGi-mCherry) at −1.5mm posterior to bregma. **(C)** Representative image of the ZI targeted with intra-cranial infusions of DREADD-expressing mCherry viruses. **(D)** Patch-clamp recording of hSyn-hM4DGi-mCherry expressing cells in the ZI showing membrane hyperpolarization during CNO exposure. **(E)** Chemogenetic inhibition of the ZI (hM4DGi+CNO) resulted in a significant increase in fear response to CS-compared to controls (GFP+CNO). **(F)** Chemogenetic inhibition of the ZI (hM4DGi+CNO) resulted in an impaired ability to discriminate between the CS+ and the CS-. \*\**p*<0.01, \*\*\*\**p*<0.0001. Data represented as Mean ± S.E.M.

### Histology

To validate the placement of intra-cranial virus injections, animals were anesthetized and trans-cardially perfused after behavioral experiments with 4% paraformaldehyde dissolved in phosphate-buffered saline (PBS). Brains were removed and stored in paraformaldehyde solution for a day and transferred to 30% sucrose solution for 3-4 days before sectioning on a freezing microtome (Leica). Brains were sectioned at 35μm, stained with Hoechst nuclear stain (1:1000) and mounted on slides using SlowFade Gold Antifade mountant (Life Technologies). The position of GFP or mCherry positive cells was assessed using Nikon Eclipse E800 fluorescent microscope.

### Open field test

The open field arena (50 × 50 × 50 cm3) was illuminated by red lights with the center defined as 16% of the total area. The mice were acclimated to the red-light conditions in the testing room for 1 hr after i.p. CNO injections (1mg/kg). The mice were then placed in the center of the arena and allowed to explore for 5 mins. Each session was videotaped using an overhead digital camera and the data was analyzed using automated video tracking system TopScan 2.0 (CleverSys Inc.).

### Electrophysiology

4-6 weeks after viral injections, 300 µm mouse brain slices containing ZI were obtained as previously reported (37). Briefly, each mouse in this study was anesthetized with isoflurane, the brain was quickly removed from the skull, and a tissue block containing the ZI mounted on the stage of a Leica VTS-1000 vibratome (Leica Microsystems Inc., Bannockburn, IL, USA). Coronal slices were obtained and then incubated in 95%O2/5%CO2 oxygenated artificial cerebrospinal fluid (ACSF) at 32°C for 1 hr before recording.

At the start of each recording, an individual slice was transferred to a recording chamber mounted on the stage of Leica STP6000 microscope and perfused with oxygenated ACSF at 32°C at a speed of 1-2 ml/min. Individual neurons in the ZI were visualized in bright field space using an infrared sensitive Hamamatsu CCD camera connected to a Windows PC using Simple PCI software. To identify neurons expressing the fluorescent transgene, we used epifluorescent illumination in combination with the appropriate excitation/emission filter sets. Standard whole cell patch-clamp recordings from fluorescent neurons in the ZI were performed using a MultiClamp 700B amplifier, an Axon Digidata 1550 A-D interface, and pClamp 10.4 software (Molecular Devices Corporation, Sunnyvale, CA). Recording pipettes were pulled from borosilicate glass and had resistances of 4-6 MΩ when filled with intracellular solution (in mM): 130 K-Gluconate, 2 KCl, 10 HEPES, 3 MgCl2, 5 phosphocreatine, 2 K-ATP, and 0.2 NaGTP. The patch solution was buffered to a pH of 7.3 and had an osmolarity of 280-290 mOsm. Current clamp recordings were performed to examine the effect of bath application of clozapine-N-oxide (CNO, 20 µM) on the resting membrane potential, and basic physiological properties of ZI neurons.

### Statistical Analysis

GraphPad Prism was used to analyze the data. Unpaired t-tests were used for data sets containing only two groups and one dependent variable (C-FOS immunohistochemistry). Repeated-measures two-way ANOVA was used to analyze data sets with more than one independent variable (behavior experiments). Post-hoc tests were only performed when interaction effects between the independent variables were significant and Sidak’s correction applied to account for multiple comparisons. Significance was set at p < 0.05.

## RESULTS

### Decreased neuronal activity in the ZI accompanies fear generalization that manifests after conditioning with high intensity foot-shocks

We trained mice in a differential auditory fear conditioning protocol using low and high threat conditions to study the role of the zona incerta in fear generalization (Fig. 1A). Wild type mice trained under low threat conditions (0.3mA foot-shocks), exhibited appropriate fear responses as indicated by increased freezing to CS+ (conditioned auditory stimulus) and reduced freezing to CS-(neutral auditory stimulus) (Fig. 1B). Under high threat conditions (0.8mA foot-shocks), wild type mice exhibited enhanced fear generalization as indicated by increased freezing to both the CS+ and CS-tones (Fig. 1B). (Low-intensity training group n = 14, High-intensity training group n = 10, Training x Tone interaction F (1, 22) = 19.17, p < 0.0001 Post-hoc tests: Low Intensity Training: CS-vs. Low Intensity Training: CS+ p<0.0001, Low Intensity Training: CS-vs. High Intensity Training: CS-p<0.01). Animals trained under high threat conditions showed poor discrimination in their fear response to the CS+ and CS- and increased generalization, as noted by their lower discrimination index compared to animals trained under low threat conditions. (Fig. 1C) (p < 0.01, t = 3.640, df = 22). Importantly, there was no statistically significant difference between the groups in their freezing response to the context alone on the day of testing (Supplementary Fig. 1), demonstrating a specificity of freezing responses to the tones.

To examine neuronal activation of the ZI in the context of fear generalization, we counted the number of cells expressing the immediate early gene, C-FOS in the ZI after exposing animals to either CS- or CS+ tone presentations. These animals had been previously trained under low threat or high threat conditions as outlined (Supplementary Fig. 2A). Animals trained under high threat conditions expressed increased fear to CS- on the day of testing, accompanied by lower numbers of C-FOS positive cells in the ZI (Figs. 1D-F, Supplementary Fig. 2B). We did not find any significant differences between groups in the numbers of C-FOS positive cells in the ZI after exposure to the CS+. (Training x Tone interaction F (1, 19) = 4.944, p < 0.05. Post-hoc tests: Low Intensity Training: CS- vs. High Intensity Training: CS- p<0.01. CS-: Low-intensity shock group n = 7, High-intensity shock group n = 8; CS+: Low-intensity shock group n = 4, High-intensity shock group n = 4). We found that in general, higher levels of fear expression (as measured by the freezing responses) were associated with lower numbers of C-FOS expressing cells in the ZI (Fig. 1G) (n = 21 animals, p < 0.01, r = −0.5563).

### Decreasing cellular activity in the ZI results in fear generalization after conditioning with low intensity foot-shocks

We utilized Gi-coupled DREADDs (Designer Receptors Exclusively Activated by Designer Drugs) to decrease activity of cells in the ZI (Fig. 2A-2D). We queried whether reducing cellular activity in the ZI would facilitate fear generalization in animals trained under low threat conditions that normally do not exhibit fear generalization. We injected AAV5-hsyn-hM4D(Gi)-mCherry or AAV5-hsyn-GFP bilaterally into the ZI of wild type mice and CNO was administered intraperitoneally, one hour before testing fear generalization. Decreasing activity of the ZI resulted in fear generalization in animals trained under low threat conditions (Fig. 2E). Specifically, Low Intensity Training-hM4D(Gi)+CNO animals exhibited significantly higher freezing responses to CS- than compared to freezing responses to the CS- of the Low Intensity Training-GFP+CNO animals. (Low Intensity Training-GFP+CNO group n = 6, Low Intensity Training-hM4DGi+CNO n = 7, DREADD x Tone interaction F (1,11) = 6.335, p < 0.05. Post-hocs: Low Intensity Training-GFP+CNO:CS- vs. Low Intensity Training-GFP+CNO:CS+ p < 0.0001, Low Intensity Training-hM4DGi+CNO:CS- vs. Low Intensity Training- hM4DGi+CNO:CS+ p < 0.01, Low Intensity Training-GFP+CNO:CS- vs. Low Intensity Training- hM4DGi+CNO:CS- p < 0.0001). Low Intensity Training-hM4DGi+CNO animals showed an impaired ability to discriminate between the CS+ and CS- as noted by their lower discrimination index compared to Low Intensity Training-GFP+CNO animals (Fig. 2F) (p < 0.01, t=3.572 df=14). There was no statistically significant difference between the groups in their freezing responses to the context (Context B) before tone presentations on the day of testing (Supplementary Fig. 3), suggesting a specificity of freezing responses to the tones. Chemogenetic inhibition of cells in the ZI was not accompanied by alterations in locomotor activity or anxiety-like behavior (Supplementary Fig. 4).

### Increasing cellular activity in the ZI reduces fear generalization that manifests after conditioning with high intensity foot-shocks

We utilized Gq-coupled DREADDs (Designer Receptors Exclusively Activated by Designer Drugs) to increase activity of cells in the ZI (Fig. 3A-3C). We queried whether stimulating cells in the ZI can reduce fear generalization observed in animals trained under high threat conditions. We injected AAV5-hsyn-hM3D(Gq)-mCherry or AAV5-hsyn-GFP bilaterally into the ZI of wild type mice and CNO was administered intraperitoneally, one hour before testing fear generalization. Increasing activity of the ZI reduced fear generalization in animals trained under high threat conditions (Fig. 3D). Specifically, High Intensity Training-hM3D(Gq)+CNO animals exhibited significantly lower freezing responses to CS- than their responses to CS+, compared to the High Intensity Training-GFP+CNO animals. (High Intensity Training-GFP+CNO group n = 7, High Intensity Training-hM3DGq+CNO n = 10, DREADD x Tone interaction F (1,15) = 20.16, p < 0.001. Post-hocs: High Intensity Training-GFP+CNO:CS- vs. High Intensity Training- hM3DGq+CNO:CS- p<0.0001, High Intensity Training-hM3DGq+CNO:CS+ vs. High Intensity Training-hM3DGq+CNO:CS- p<0.0001, High Intensity Training-GFP+CNO:CS+ vs. High Intensity Training-hM3DGq+CNO:CS+ p<0.01). High Intensity Training-hM3DGq+CNO animals showed better discrimination in their fear response to the CS+ and CS- as noted by their higher discrimination index compared to High Intensity Training-GFP+CNO animals (Fig. 3E) (p < 0.0001, t = 5.931, df = 17). There was no statistically significant difference between the groups in their freezing responses to the context (Context B) before tone presentations on the day of testing (Supplementary Fig. 5), suggesting a specificity of freezing responses to the tones. These observed differences in freezing responses of animals with chemogenetic activation of ZI, were not accompanied by alterations in locomotor activity or anxiety-like behavior (Supplementary Fig. 6).

**Figure 3:**
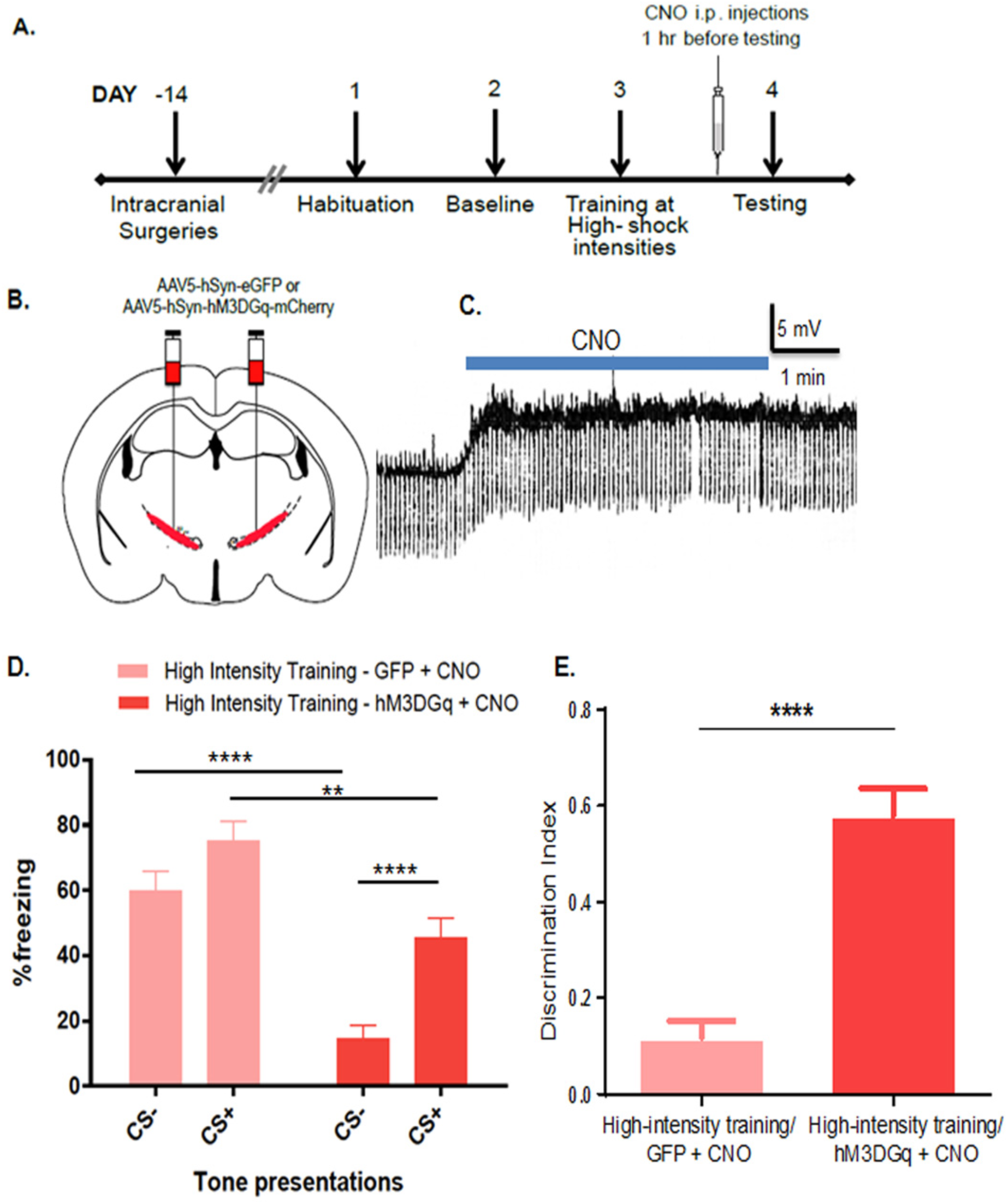
Increasing cellular activity in the ZI prevents fear generalization. **(A)** Experimental protocol for chemogenetic activation. Two weeks after intracranial injection of the control or DREADD virus, animals were conditioned to high threat intensities. The next day, CNO was administered intraperitoneally 1 hour before testing fear generalization. **(B)** Wild-type animals were injected with either the control virus (AAV5-hSyn-eGFP) or excitatory DREADDs (AAV5-hSyn-hM3DGq-mCherry) at −1.5mm posterior to bregma. **(C)** Patch-clamp recording of hSyn-hM3DGq-mCherry expressing cells in the ZI showing membrane depolarization during CNO exposure. **(D)** Training using high intensity foot-shock causes fear generalization as seen by high freezing to both CS+ and CS-. Chemogenetic activation of the ZI (hM3DGq+CNO) resulted in a significant decrease in fear response to CS+ as well as CS- compared to controls (GFP+CNO). **(E)** Chemogenetic activation of the ZI (hM3DGq+CNO) resulted in a better ability to discriminate between the CS+ and the CS-. \*\**p*<0.01, \*\*\*\**p*<0.0001. Data represented as Mean ± S.E.M.

### Increasing activity of GABAergic cells in the ZI reduces fear generalization that manifests after conditioning with high intensity foot-shocks

GABAergic cells in the ZI have been implicated in defensive responses like freezing and avoidance as well as in retrieval of aversive memories (29, 30). We tested whether targeted stimulation GABAergic cells in the ZI can reduce fear generalization (Fig. 4A). We injected AAV5-DIO-hSyn-hM3D(Gq)-mCherry or AAV5-DIO-hSyn-mCherry bilaterally into the ZI of vGAT-CRE mice (Figs. 4B, 4C). CNO was administered intraperitoneally, one hour before testing fear generalization. Increasing activity of GABAergic cells in the ZI alone, drastically reduced fear generalization observed in animals trained under high threat conditions (Fig. 4D). Specifically, vGAT-CRE:DIO-hM3D(Gq)-mCherry+CNO animals exhibited significantly lower freezing responses to CS- than their responses to CS+, compared to vGAT-CRE:DIO-mCherry+CNO animals. (vGAT-CRE:DIO-mCherry+CNO group n = 9, vGAT-CRE:DIO-hM3D(Gq)-mCherry+CNO n = 11, DREADD x Tone interaction F (1,18) = 21.48, p < 0.0001. Post-hocs: High Intensity Training-DIO-hM3DGq+CNO:CS+ vs. High Intensity Training-DIO-hM3DGq+CNO:CS- p<0.0001, High Intensity Training-DIO-GFP+CNO:CS- vs. High Intensity Training-DIO-hM3DGq+CNO:CS- p<0.0001, High Intensity Training-DIO-GFP+CNO:CS+ vs. High Intensity Training-DIO-hM3DGq+CNO:CS+ p<0.05). High Intensity Training-DIO-hM3DGq+CNO animals showed better discrimination in their fear response to the CS+ and CS- as noted by their higher discrimination index compared to High Intensity Training-DIO-GFP+CNO animals (Fig. 4E) (p < 0.0001, t = 9.151, df = 18). There was no statistically significant difference between the vGAT-CRE groups in their freezing to the context (Context B) before tone presentations on the day of testing (Supplementary Fig. 7), suggesting a specificity of freezing responses to the tones. These observed differences in freezing responses of animals with chemogenetic stimulation of GABAergic cells, were not accompanied by alterations in locomotor activity or anxiety-like behavior (Supplementary Fig. 8).

**Figure 4:**
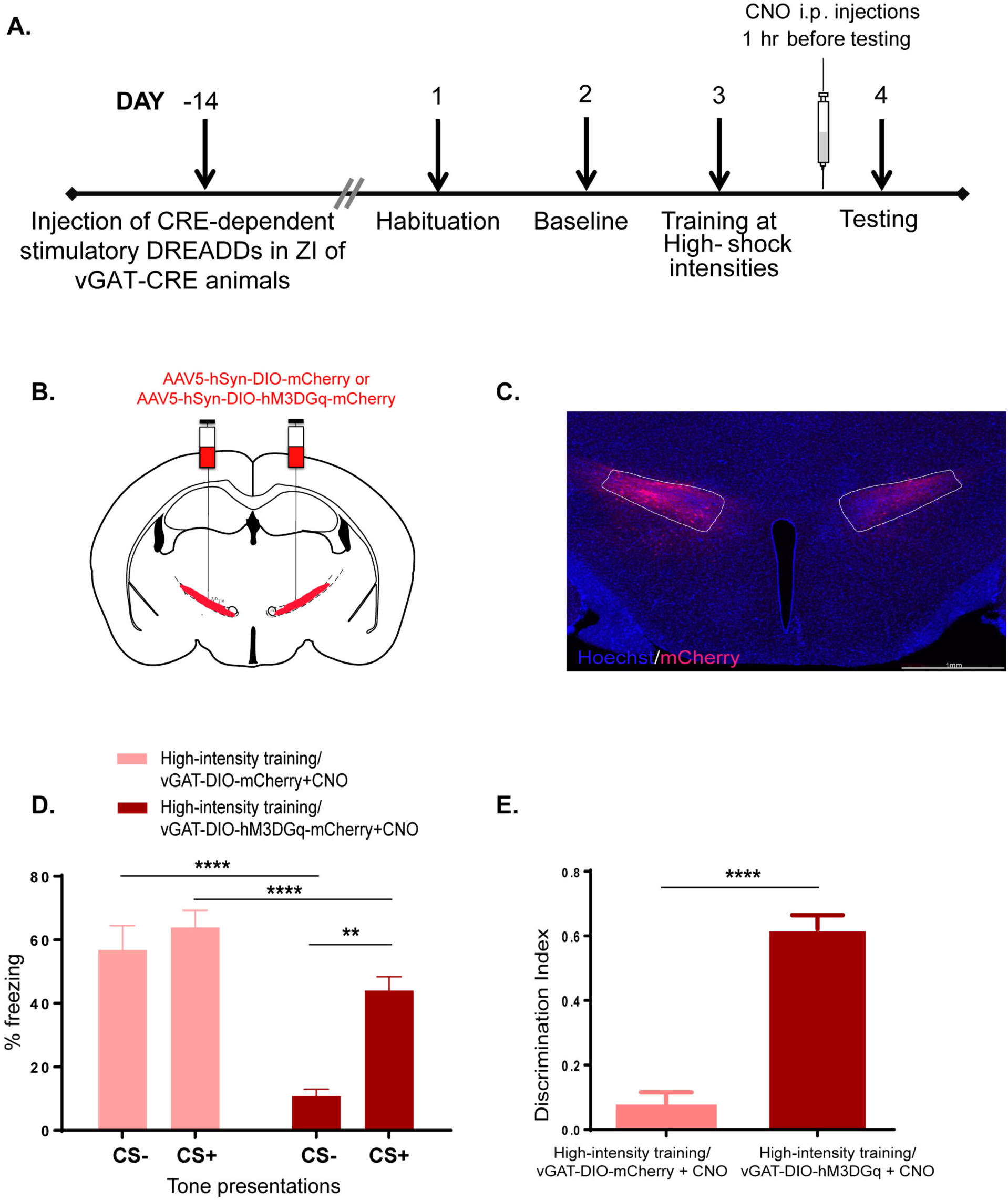
Targeted chemogenetic activation of GABAergic cells in the ZI can reduce fear generalization. **(A)** Experimental design: vGAT-CRE animals received intracranial injections of CRE-dependent control or DREADD virus and after two weeks, were conditioned to high threat intensities. One day post-training, CNO was administered intraperitoneally 1 hour before testing for fear generalization. **(B)** vGAT-CRE animals were injected with either the control virus (AAV5- hSyn-DIO-mCherry) or excitatory DREADDs (AAV5-hSyn-DIO-hM3DGq-mCherry) at −1.5mm posterior to bregma. **(C)** Representative image of the GABAergic cells within the ZI infected with mCherry-expressing excitatory DREADDs. **(D)** Animals with expression of DIO-hM3DGq virus in vGAT-CRE expressing GABAergic cells in the ZI and injected with CNO (DIO-hM3DGq+CNO) one hour before testing for fear generalization showed a significant decrease in fear response to CS- compared to animals that were infused with the DIO-mCherry virus in vGAT-CRE expressing GABAergic cells in the ZI and injected with CNO (DIO-mCherry+CNO). **(E)** Chemogenetic activation of GABAergic cells in the ZI (DIO-hM3DGq+CNO) resulted in a better ability to discriminate between the CS+ and the CS-. \**p*<0.05 **** *p*<0.0001. Data represented as Mean ± S.E.M.

## DISCUSSION

Our results demonstrate a novel role for the zona incerta (ZI) in modulating fear generalization. First, we found reduced C-FOS activation in the ZI associated with increased fear towards a neutral auditory stimulus and that reducing cellular activity in the ZI resulted in fear generalization in animals trained even under low threat conditions. Next, we found that stimulating cellular activity in the ZI using a chemogenetic strategy reduced generalized fear responses observed after training animals under high threat conditions. Finally, we demonstrated that targeted stimulation of GABAergic cells in the ZI reduced fear generalization resulting from exposure to high threat training conditions. Taken together, our data provide evidence for a translationally relevant role for the ZI in modulating fear generalization.

Lesioning studies as well as computational models have suggested a role for thalamic and sub-thalamic brain circuits in stimulus discrimination (38-40) – a key component of fear generalization. In particular, it is hypothesized that the broader receptive fields of thalamic and sub-thalamic neurons communicating with core fear-related circuitry could support fear generalization (39, 41, 42). Increasing threat intensities broaden generalization gradients in humans (43). In this study, we used a translationally relevant threat intensity-based model of fear generalization in rodents to examine sub-thalamic contributions to fear processing. Animals when trained under high threat conditions (0.8mA foot-shocks) generalized fear to both conditioned (CS+) and neutral (CS-) tones whereas animals trained under low threat conditions (0.3mA foot-shocks) did not demonstrate such generalization. These observations agree with previous reports that conditioning using increasing shock intensities promotes fear generalization in rodents (8, 44).

To test whether the ZI is responsive to neutral stimuli and potentially involved in fear generalization, we first sought to compare C-FOS immunohistochemistry in the ZI of animals trained under low and high threat conditions. More specifically, we counted C-FOS positive cells in the ZI of animals exposed to the CS+ or CS- on testing day. Excitingly, we found fewer C-FOS positive cells in the ZI of animals that generalized fear to the CS-. Additionally, we found reduced C-FOS expression in the ZI after exposure to the CS+ in animals trained under low as well as high threat conditions. Could the ZI modulate fear generalization associated with high threat conditions? The ZI is ideally positioned to convey information regarding the salience of specific sensory stimuli and orchestrate appropriate fear-related behavioral responses. First, the ZI receives projections from sensory cortices (including the auditory cortex) and can coordinate activity across cortical networks according to attentional demands (45, 46). Additionally, the ZI innervates midbrain regions like the periaqueductal gray that plays an important role in orchestrating fearful behaviors (18, 47, 48). Second, the ZI has been implicated in sensory discrimination and can modulate incoming sensory information (49). Finally, stimulating the ZI in humans facilitates the discrimination of fearful faces from non-fearful ones (33, 50). It is possible that stressful states like those created by high-intensity threat conditioning directly perturb cellular function in the ZI, rendering fear generalization as a behavioral outcome. Alternatively, generalized fear responses could arise indirectly due to amygdala→ZI connectivity (30, 47). Loss of cue-specificity and widening of the memory trace in the amygdala occurring during fear generalization (8) could alter ZI’s influence on modulating fear responses. Future experiments will need to examine how cellular and molecular niches in the ZI are impacted by stress as well as amygdala function, resulting in generalization of fear responses.

Building on our observations from the C-FOS study, we used DREADD-based strategies to test whether manipulating cellular activity in the ZI affected fear generalization. Reducing cellular activity in the ZI resulted in animals trained under low threat conditions showing fear generalization. Conversely, increasing the activity of cells in the ZI of animals trained under high threat conditions distinctly reduced fear generalization. Given the complex neurochemical profile of ZI (51), we wanted to determine the cell populations responsible for suppressing fear generalization. The GABAergic neurons of the ventral ZI were of particular interest, since they have been shown to gate ascending sensory information by fast feed-forward inhibition of higher order thalamic nuclei (22, 49, 52, 53). More recently, GABAergic cells in the ZI have been shown to be important for defensive responses as well as acquisition and retrieval of fear memories (29, 30). In our study, we found that stimulating GABAergic cells in the ZI reduced fear generalization in animals conditioned under high threat intensities. It is important to note that the caudal ZI has been associated with motor function due to its connections with the basal ganglia network and has been investigated as a potential target for deep brain stimulation treatment for patients with Parkinson’s disease (PD) (33, 47, 50). Therefore, alterations in locomotor behavior could have potentially contributed to the observed effects on fear generalization following the bidirectional chemogenetic manipulations of cellular activity in the ZI. However, we did not observe any significant differences in total distance traveled and velocity during an open field test performed after chemogenetic stimulation of ZI (Supplementary Figs. 4A, 4B, 6A, 6B, 8A, 8B). Freezing to the testing context (Context B) remained unaltered after stimulating activity in the ZI (Supplementary Figs. 3, 5, 7), further emphasizing that the observed effects on fear generalization were specific to the CS+ and CS- tones presented.

Stimulating GABAergic cells and the paravalbumin (PV)-expressing cells in the ZI has been demonstrated to reduce fearful behavior (29, 30). In line with these findings, we find that stimulation of GABAergic cells within the ZI results in reduced fear responses toward the conditioned stimuli. A novel component of our study is the finding that stimulating GABAergic cells in the ZI enhances discrimination between the CS+ and CS- and reduces generalization as illustrated by the altered discrimination index and reduced fear response to the neutral stimulus. In so doing, stimulating the ZI still leaves room for adaptive fear responses to the CS+ to be expressed while reducing fear to the neutral CS-.

As noted above, the ZI is chemo-architecturally diverse and future experiments will be needed to determine specific neuromodulators (e.g. parvalbumin or somatostatin or both) present in the GABAergic cells in the ZI and the role of their downstream targets in tuning generalization of fear responses. Blocking synaptic transmission in the ZI has also been shown to alter anxiety-related measures (30). However, we did not find similar effects with chemogenetic manipulation of the ZI (Supplementary Figs. 4C, 6C, 8C). These stated differences emphasize the importance of dissecting contributions of specific cell-types in the ZI to varied dimensions of fear and anxiety.

Overall, the experimental results described here bolster the recently demonstrated link between ZI activity and fearful behavior and its role in calibrating fearful behavior toward environmental stimuli. Our study makes a novel contribution to this body of work by demonstrating a role for the ZI in fear generalization. To conclude, our work suggests that stimulating the ZI in the clinic during exposure therapy could reduce fear generalization, while leaving adaptive fear responses intact.

## Acknowledgements

We thank the Veterinary and Animal Care staff in the Yerkes Neuroscience Vivarium for animal husbandry. Funding for this study was provided to BGD by the Department of Psychiatry and Behavioral Sciences, the Brain Health Institute, the Yerkes National Primate Research Center (YNPRC), a CIFAR Azrieli Global Scholar award, and the Catherine Shopshire Hardman Fund. Additional funding was provided to YNPRC by Office of Research Infrastructure Programs ODP51OD11132.

## Author Contributions

AV and BGD conceptualized the project. AV, and BGD designed the study. AV, NB, PR, JG and BGD performed experiments, analyzed data, and interpreted results. AV, JG and DR helped with manuscript preparation. AV and BGD wrote the manuscript.

## Financial Disclosures/Conflict of Interest

The authors have no competing financial interests or potential conflicts of interest.

---

**Supplementary Figure 1:**
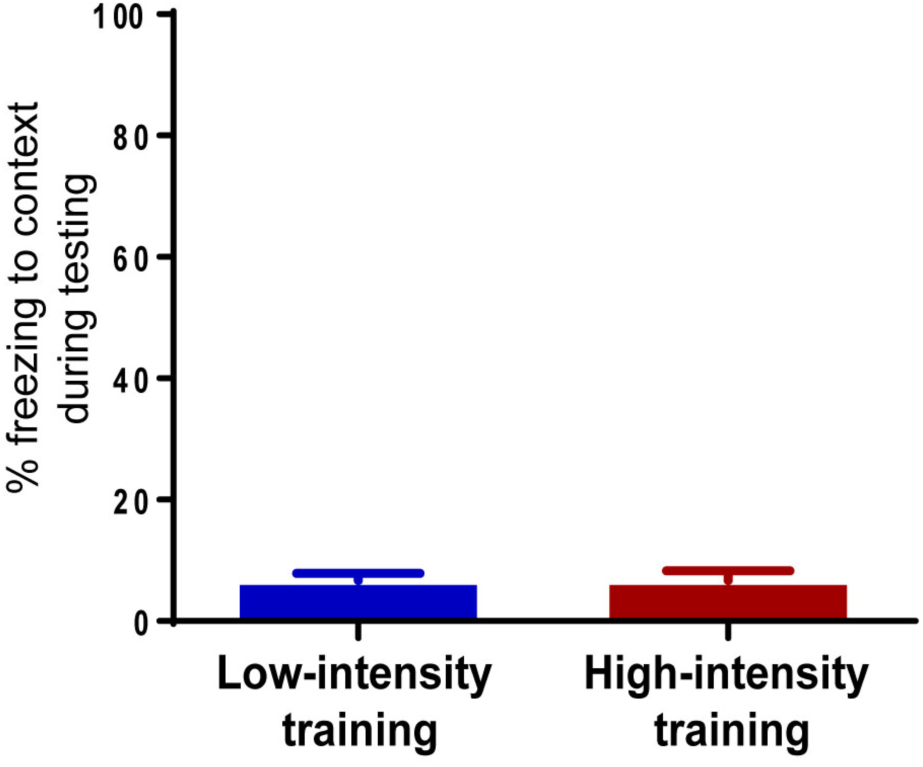
Freezing responses in both groups are specific to the tone presentations on testing day. No significant differences were observed in freezing to Context B on testing day between animals trained under low and high threat intensities. This demonstrates that the observed differences in fear generalization is specific to the tone presentations and does not indicate an overall change in fear response in the animals. Data represented as Mean ± S.E.M.

**Supplementary figure 2:**
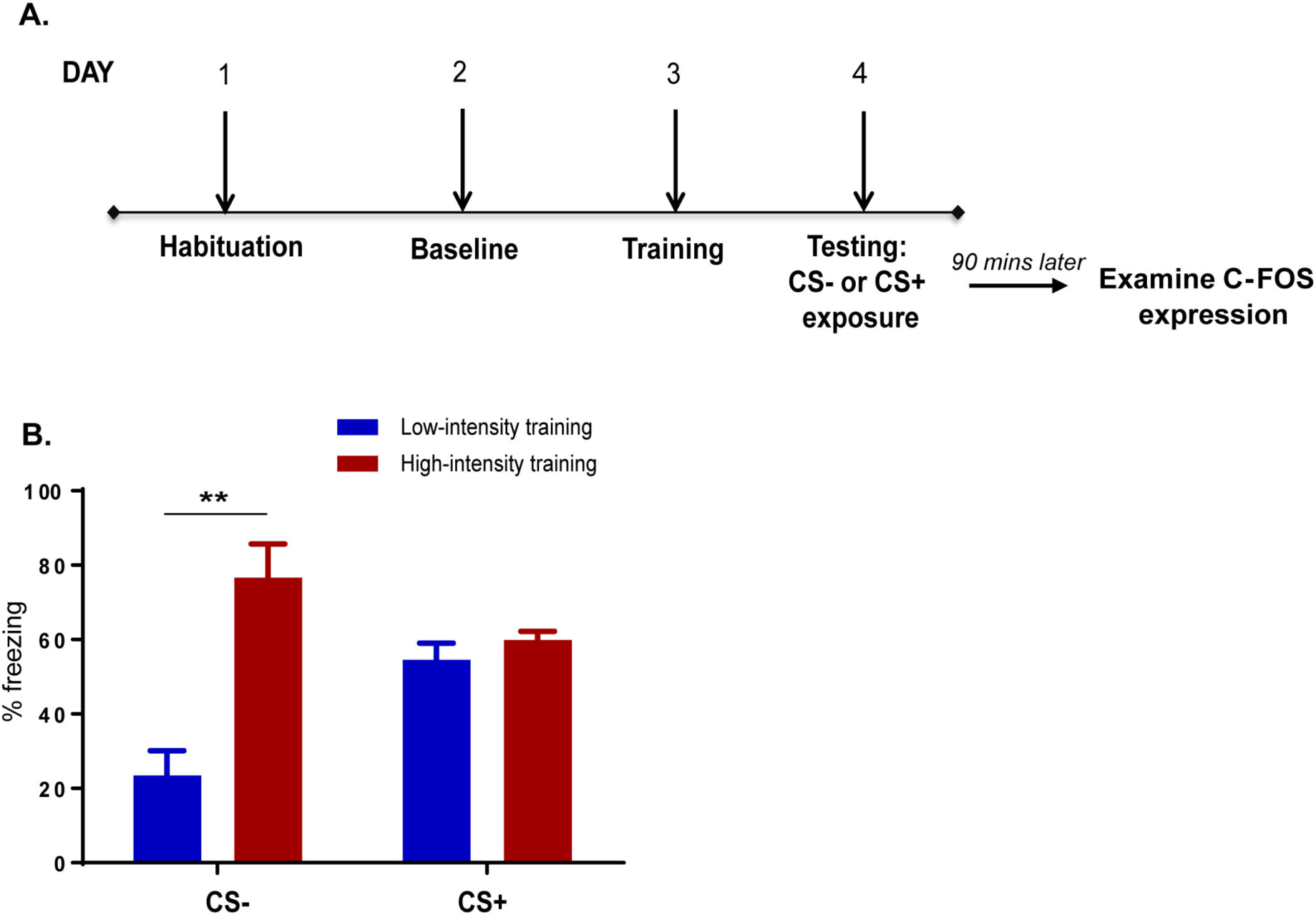
Animals trained under high threat conditions express increased fear to the neutral stimulus alone. **(A)** Experimental design for C-FOS study: After habituation and baseline recording of stimulus responses, animals were split into four different groups. One group of mice received CS+ tone presentations paired with 0.3mA foot-shocks (Low-intensity training) and unpaired CS-tone presentations. Another group of mice received CS+ tone presentations paired with 0.8mA foot-shocks (High-intensity training) and unpaired CS-tone presentations. On day 2, each group was further divided in to two and freezing responses were recorded for CS+ or CS-tone presentations alone (Low-intensity training/CS-; Low-intensity training/CS+; High-intensity training/CS-; High-intensity training/CS+). **(B)** Animals trained under high threat conditions show significantly increased freezing response to CS+ compared to animals trained under low threat conditions. No significant differences were observed in animals’ response to CS+, when trained under the different threat intensities.

**Supplementary Figure 3:**
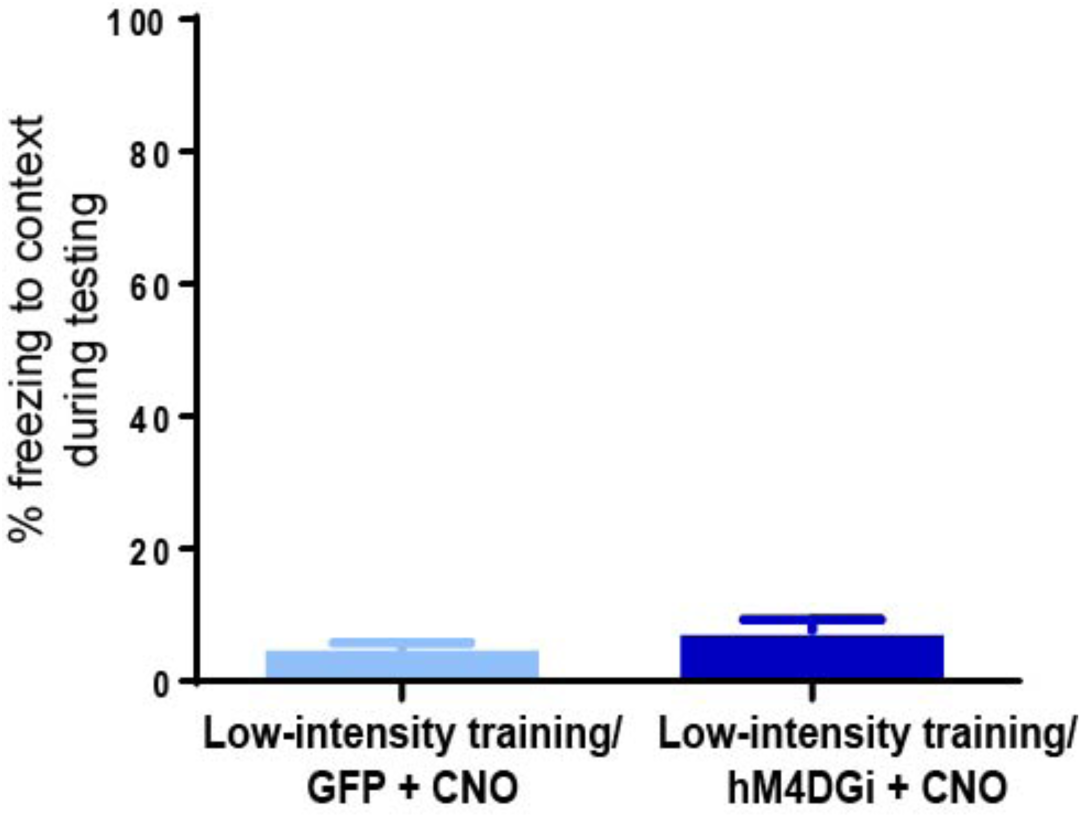
Chemogenetic inhibition of the ZI does not produce non-specific alterations in freezing responses. No significant differences were observed in freezing to Context B with chemogenetic inhibition of the ZI on testing day, demonstrating that the observed changes in freezing responses (Fig. 2) were specific to the auditory stimuli. Data represented as Mean ± S.E.M.

**Supplementary Figure 4:**
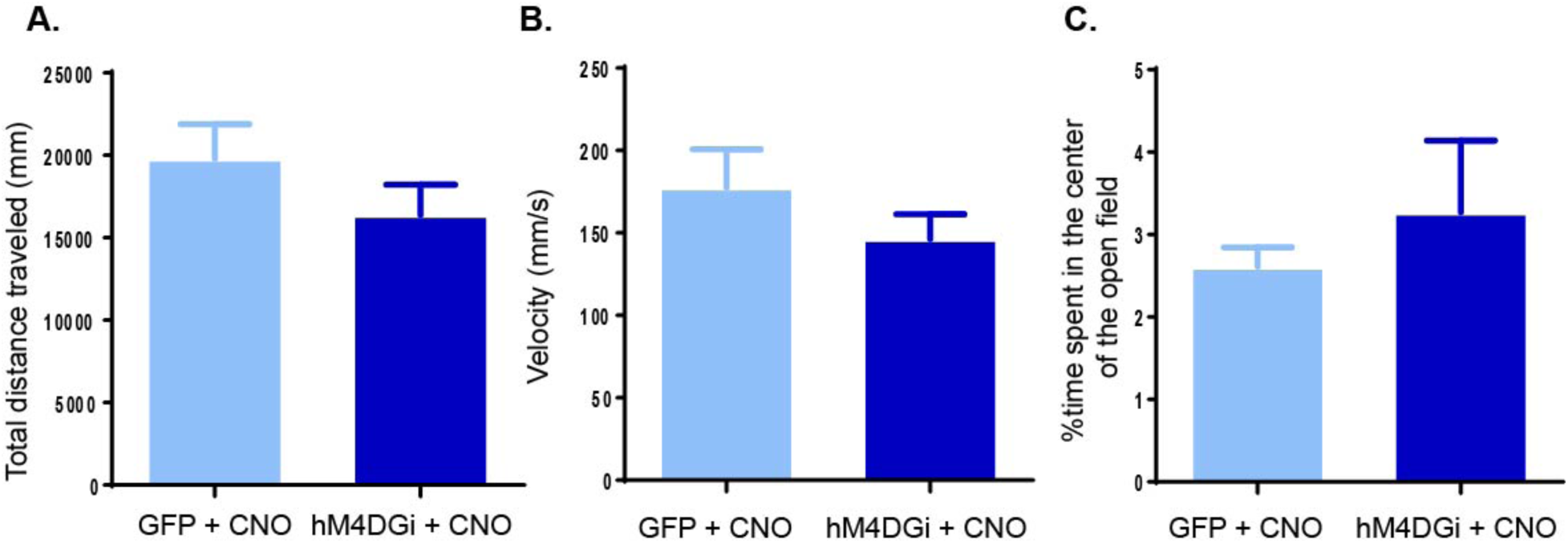
Chemogenetic inhibition of the ZI does not affect general locomotor function and anxiety-like behavior. In the Open Field Test performed one hour after CNO injections, chemogenetic inhibition (hM4DGi+CNO) of the ZI in wild type animals did not produce detectable changes in **(A)** total distance traveled (in mm), **(B)** velocity (mm/sec), and **(C)** time spent in center of open field, compared to controls (GFP+CNO). Data represented as Mean ± S.E.M.

**Supplementary Figure 5:**
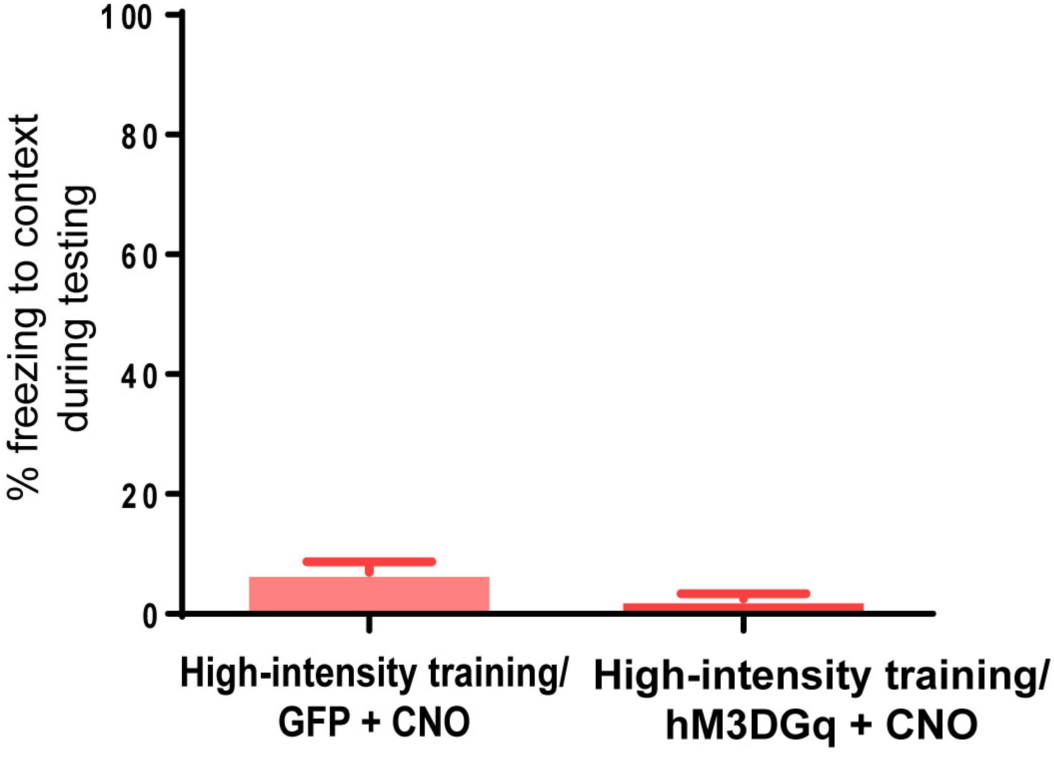
Chemogenetic activation of the ZI does not produce non-specific increase in freezing responses. No significant differences were observed in freezing to Context B with chemogenetic activation of ZI on testing day, demonstrating that the observed changes in freezing responses (Fig. 3) were specific to the auditory stimuli. Data represented as Mean ± S.E.M.

**Supplementary Figure 6:**
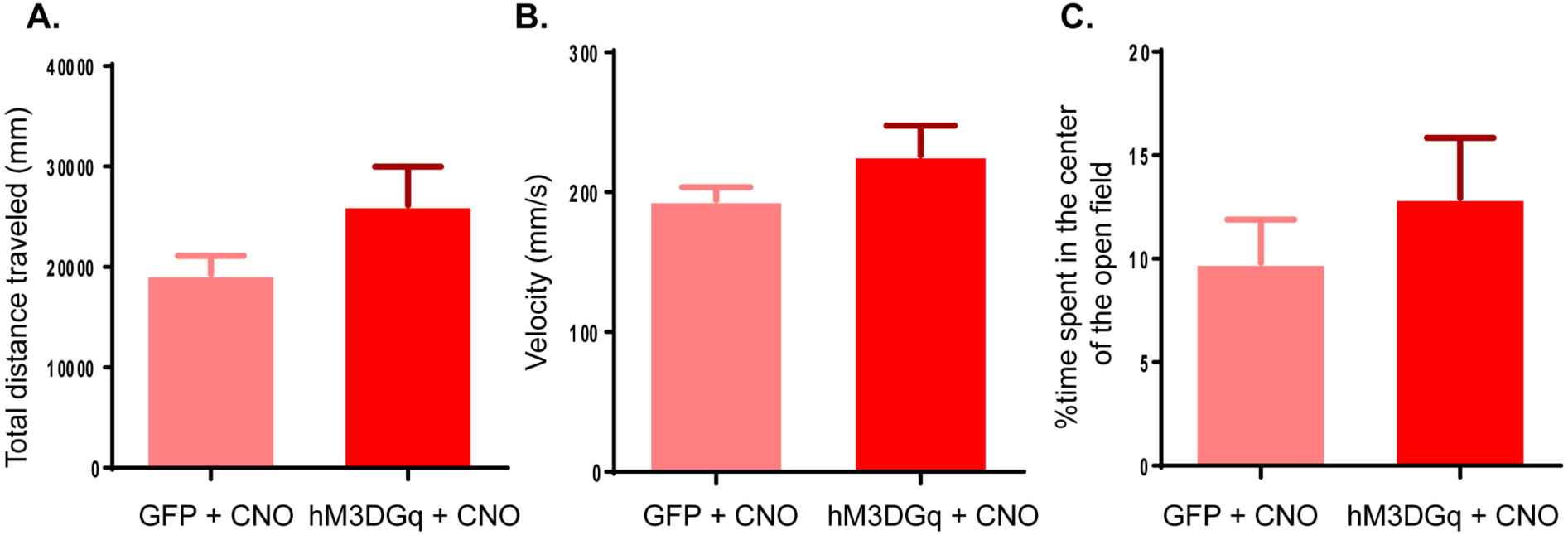
Chemogenetic activation of the ZI does not affect general locomotor function and anxiety-like behavior. In the Open Field Test performed one hour after CNO injections, chemogenetic activation (hM3DGq+CNO) of the ZI in wild type animals did not produce detectable changes in **(A)** total distance traveled (in mm), **(B)** velocity (mm/sec), and **(C)** time spent in center of open field, compared to controls (GFP+CNO). Data represented as Mean ± S.E.M.

**Supplementary Figure 7:**
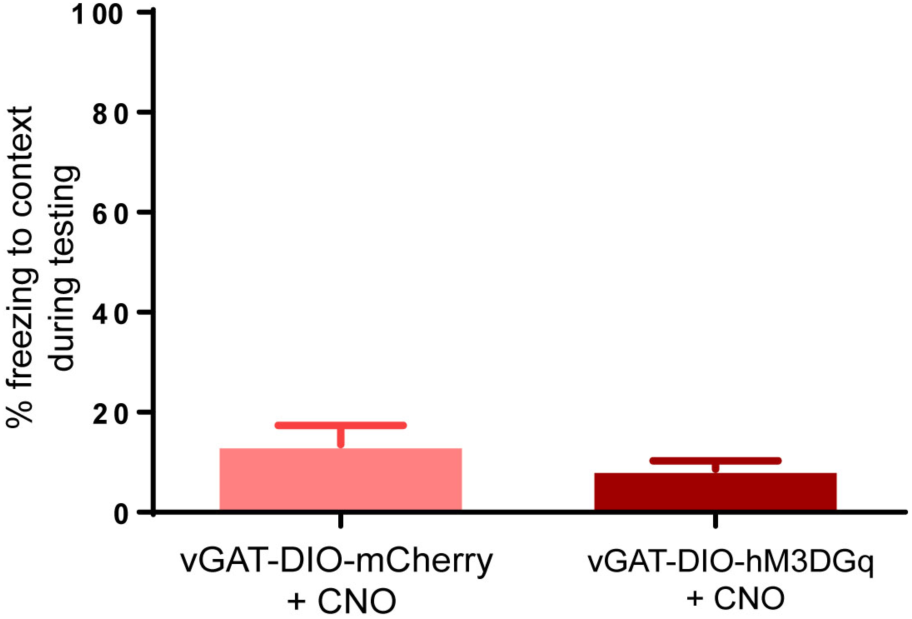
Chemogenetic activation of GABAergic cells in the ZI does not produce non-specific increase in freezing responses. **(A)** No significant differences were observed in freezing to Context B with chemogenetic activation of GABAergic cells in the ZI on testing day. These data show that the observed changes in freezing responses following chemogenetic stimulation of GABAergic cells in the ZI (Fig. 4) were specific to the auditory stimuli. Data represented as Mean ± S.E.M.

**Supplementary Figure 8:**
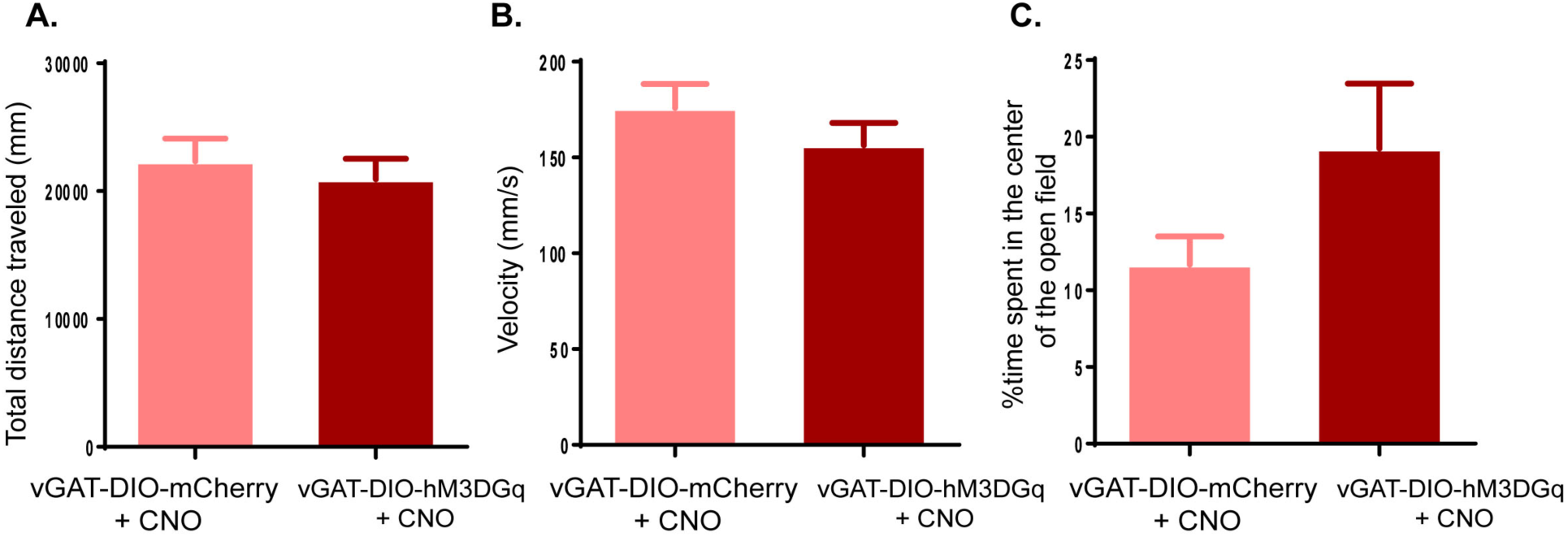
Chemogenetic activation of GABAergic cells in the ZI does not affect general locomotor function and anxiety. In the Open Field Test performed one hour after CNO injections, chemogenetic activation (DIO-hM3DGq+CNO) of GABAergic cells in the ZI in vGAT-CRE animals did not produce detectable changes in **(A)** total distance traveled (in mm), **(B)** velocity (mm/sec), and **(C)** time spent in center of open field, compared to controls (DIO-GFP+CNO). Data represented as Mean ± S.E.M

